# ARGONAUTE5 Mediates Fine-Tuning of Vegetative-to-Reproductive Phase Transition Through Its Interaction with miR156 in *Arabidopsis*

**DOI:** 10.1101/640680

**Authors:** Charles Roussin-Léveillée, Guilherme Silva-Martins, Peter Moffett

**Affiliations:** Centre SÈVE, Département de Biologie, Université de Sherbrooke, Sherbrooke, Québec J1K 2R1, Canada

**Author notes:** These authors contributed equally to this work (Charles Roussin-Léveillée, Guilherme Silva-Martins).

**Keywords:** ARGONAUTE5, plant development, SPL, shoot apex meristem, miR156

## Abstract

Vegetative-to-reproductive phase change is a finely tuned process in plants, largely controlled by the age-regulated microRNA156 (miR156), which functions by suppressing the transcripts of SQUAMOSA-PROMOTER BINDING LIKE (SPL) transcription factors. ARGONAUTE proteins (AGO) are essential effectors of miRNA-mediated gene regulation. However, which AGO(s) mediate(s) the control of flowering time remains unclear. Here, we demonstrate a role for AGO5 in vegetative-to-reproductive phase transition through the modulation of SPL transcription factors. We show that AGO5 interacts physically and functionally with miR156 and that *ago5* mutants present an early flowering phenotype in *Arabidopsis*. Furthermore, in *ago5* mutants, the repression of flowering caused by miR156 overexpression is largely reversed, whereas leaf morphology remains unaffected. Our results thus indicate a specific role for AGO5 in mediating miR156 activity in meristematic, but not vegetative, tissue. As such, our data suggest a spatiotemporal regulation of the miR156 aging pathway, mediated through different AGO proteins in different tissues.

## INTRODUCTION

In plants, the vegetative-to-reproductive phase transition is tightly regulated by multiple genetic pathways, some of which are controlled by the expression of small non-coding RNAs (Fouracre & Poethig, 2016; Teotia & Tang, 2015; Wu et al., 2009). These include microRNAs, which are processed into 20-24 nucleotide small RNAs from endogenously encoded microRNA precursors (pri-miRNA) by DICER-LIKE (DCL) proteins and are subsequently bound by ARGONAUTE (AGO) proteins (Baulcombe, 2004; Borges & Martienssen, 2015; Weiberg & Jin, 2015). The latter proteins are guided through sequence complementarity to cleave or repress the translation of a specific mRNAs (H. Zhang, Xia, Meyers, & Walbot, 2015; Jones-Rhoades et al. 2006). Among the miRNAs regulating developmental phase transition, microRNA156 (miR156) regulates key developmental transition genes by repressing the translation of SQUAMOSA PROMOTER BINDING PROTEIN (SBP) transcription factors(He et al., 2018; Poethig, 2013; Xu et al., 2016). *Arabidopsis thaliana (Arabidopsis* hereafter) encodes for sixteen SBP-LIKE (SPL) genes, of which ten are post-transcriptionally regulated by miR156 (Gandikota et al., 2007; Rhoades et al., 2002; Schwab et al., 2005; Wu & Poethig, 2006a). Studies of shoot apex development suggest that SBP transcription factors regulate subsets of genes, such as *APETALA1* (AP1), *LEAFY* (LFY), *SUPPRESSOR OF CONSTANS1* (SOC1), *FRUITFUL*L (FUL) and *AGAMOUS-LIKE42* (AGL42) (Xu et al. 2016; Wang et al. 2009). High levels of miR156 expression can be observed early in development, gradually decreasing in expression over time, allowing the translation of multiple transition-promoting proteins (Wu & Poethig 2006; Wu et al. 2009; Wang et al. 2014).

An increase in the levels of SPL transcript abundance due to declining miR156 level drives vegetative development and vegetative-to-reproductive phase change in many plant species (Fornara & Coupland, 2009; Hibara et al., 2016; Wei et al., 2017). Previous studies have shown that miR156 targets SPL3, SPL4 and SPL5, which are involved in floral meristem identity, along with SPL2, SPL9, SPL10, SPL11 and SPL15, which control shoot maturation and floral induction (Wang et al. 2009; Schwarz et al. 2008; Xu et al., 2016). Consistent with this, overexpression of miR156 dramatically delays juvenile-to-adult and reproductive phase transition through cleavage of mRNA and translational repression of SPL proteins (Chuck, Meeley, Irish, Sakai, & Hake, 2007; Wu & Poethig, 2006a).

Although multiple reports have investigated the effects of miR156 on SPL-encoding mRNAs, less is known about which AGO proteins functionally interact with miR156 in different developmental processes. The *Arabidopsis* genome encodes for ten AGO proteins, which function in multiple regulatory mechanisms, determined at least in part by their expression patterns and by the nature of the small RNAs they bind (Vaucheret, 2008).

In *Arabidopsis*, the AGO1 and AGO10 proteins have been extensively characterized for their roles in plant development (Vaucheret, Vazquez, Crété, & Bartel, 2004; Zhu et al., 2011). AGO1 is mainly known for its importance in mediating mRNA cleavage through its interaction with miRNAs, as well as in generating trans-acting small-interfering RNA (tasiRNA), which together regulate a plethora of physiological and developmental phenomena (Baumberger & Baulcombe, 2005; Fei, Xia, & Meyers, 2013; Rogers & Chen, 2013; Vaucheret et al., 2004). Given the large number of genes regulated by AGO1, null mutants are lethal and many of the developmental phenotypes caused by hypomorphic alleles may be pleiotropic in nature. AGO10 has also been identified as a key player in meristem development through its antagonism with AGO1 by sequestering miR165/166, which negatively regulates of the HD-ZIP III gene family (Zhu et al., 2011). With respect to shoot development, AGO1 has been shown to bind miR156 in seedlings (Azevedo et al., 2010; Ronemus, Vaughn, & Martienssen, 2006) and regulates SPL2/9/11 during abiotic stress (Stief et al. 2014). However, the identity of the ARGONAUTE protein(s) that mediate(s) the effects of miR156 in vegetative-to-reproductive phase transition has not been specifically addressed.

Here, we show that *ago5* mutant plants display an early bolting phenotype, suggesting a role for AGO5 in developmental phase transition. This conclusion was supported by the observation that SBP/SPL gene expression occurred earlier in *ago5* plants. Accordingly, we found that AGO5 binds to, and stabilizes miR156. Furthermore, we show that the late flowering phenotype caused by constitutive mir156 overexpression is largely abrogated in *ago5* mutants, as are the effects of SPL and other developmental genes.

## RESULTS

### Acceleration of the Vegetative-to-Reproductive Phase Transition in *Arabidopsis ago5* mutants

The AGO5 protein is involved in antiviral defense (Brosseau and Moffett, 2015). In the course of studying this phenomenon, we first observed (regardless of virus infection) a marked difference in flowering time in two independent *Arabidopsis ago5* mutant lines. Under a long day (LD) photoperiod, we found that *ago5* mutants displayed an early vegetative-to-reproductive phase transition as compared to wild-type Columbia-0 (Col-0) (Figure 1A). Since bolting time is an important marker of vegetative-to-reproductive phase transition, we assessed the difference in this parameter between Col-0 and two *ago5* mutant lines, *ago5-1* and *ago5-5*. Bolting started on average at 23 days post-germination (dpg) in *ago5* mutants, while Col-0 bolted approximately 5 days later, at 28 dpg (Figure 1B).

**Figure 1.**
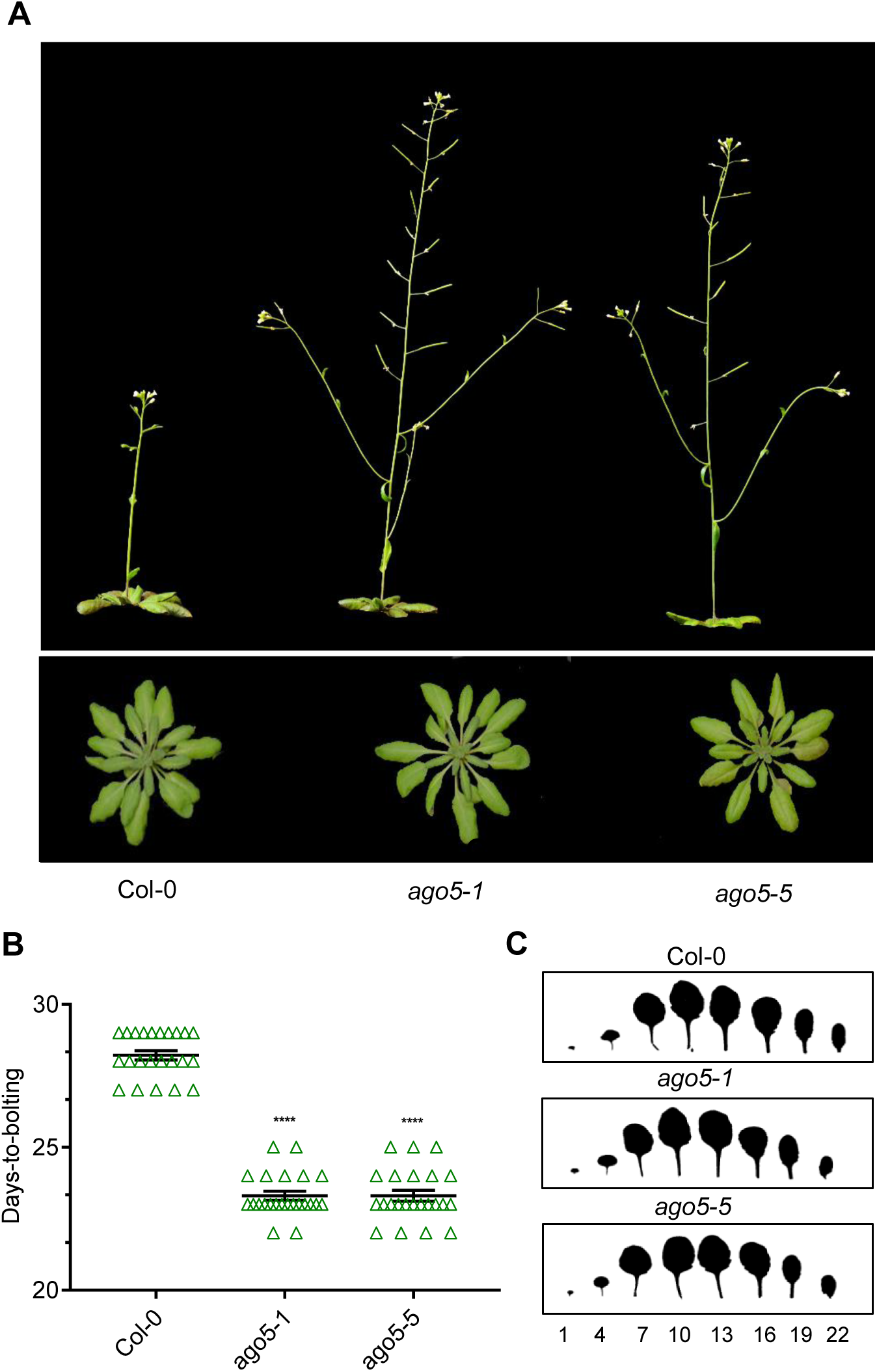
*Arabidopsis ago5* mutant plants display an early vegetative-to-reproductive phase transition phenotype. (**A**) Phenotype of Col-0 and *ago5* T-DNA insertion mutants (*ago5-1* and *ago5-5*). (**B**) Days-to-bolting of the aforementioned genotypes (n = 25). (**C**) Representative scans of fully expanded 20 days-old rosettes. Numbers represent leaves starting from the base of the rosette. Asterisks indicate significant differences from Col-0 in a Student’s t-test (*P < 0.05, * * P < 0.01, * * * P < 0.001, * * * * P < 0.0001). Error bars indicate standard deviation.

The earlier vegetative-to-reproductive transition in *ago5* mutants did not alter floral architecture or number of seeds per silique, compared to Col-0 (Figure 1 – figure supplement 1A, C). However, *ago5* mutants produced fewer flowers [138.6±2.95, 138.1±2.87 (mean±SEM) for *ago5-1* and *ago5-5*, respectively] compared to wild type [184.9±2.26 (mean±SEM)] (Figure 1 – figure supplement 1B). This difference was not due to the final height reached by the different genotypes since both *ago5* mutants and Col-0 ultimately grew to approximately the same height (data not shown).

We next investigated whether AGO5 was important in juvenile-to-adult phase transition, which differs from the vegetative-to-reproductive phase transition, and involves leaf development. However, we found no striking difference in leaf shape or morphology between Col-0 and *ago5* mutants (Figure 1C). Thus, the early bolting phenotype of *ago5* mutants is likely due to important changes in the SAM rather than a more global effect on plant maturation.

### AGO5 Is Highly Expressed in the Shoot Apical Meristem

To further study the role of AGO5 in the regulation of vegetative-to-reproductive phase transition, we investigated the levels of *AGO5* expression in different tissues. *AGO5* expression was assessed in 20 dpg plants under a LD photoperiod in both leaves and the SAM. A striking difference was found between these tissues in that *AGO5* was expressed 60-fold higher in SAM tissue compared to leaves (Figure 2A). Since control of developmental phase transition is a finely regulated phenomenon, we also assessed *AGO5* expression in the same tissues of 14 dpg plants. As expected, in 14 dpg plants *AGO5* was expressed 24-fold higher in SAM tissues, compared to leaves (Figure 2A). Interestingly, *AGO5* was 10-fold more expressed in the SAM of 20 dpg plants as compared to 14 dpg plants (Figure 2A), suggesting a developmentally-regulated progression in *AGO5* expression levels.

**Figure 2.**
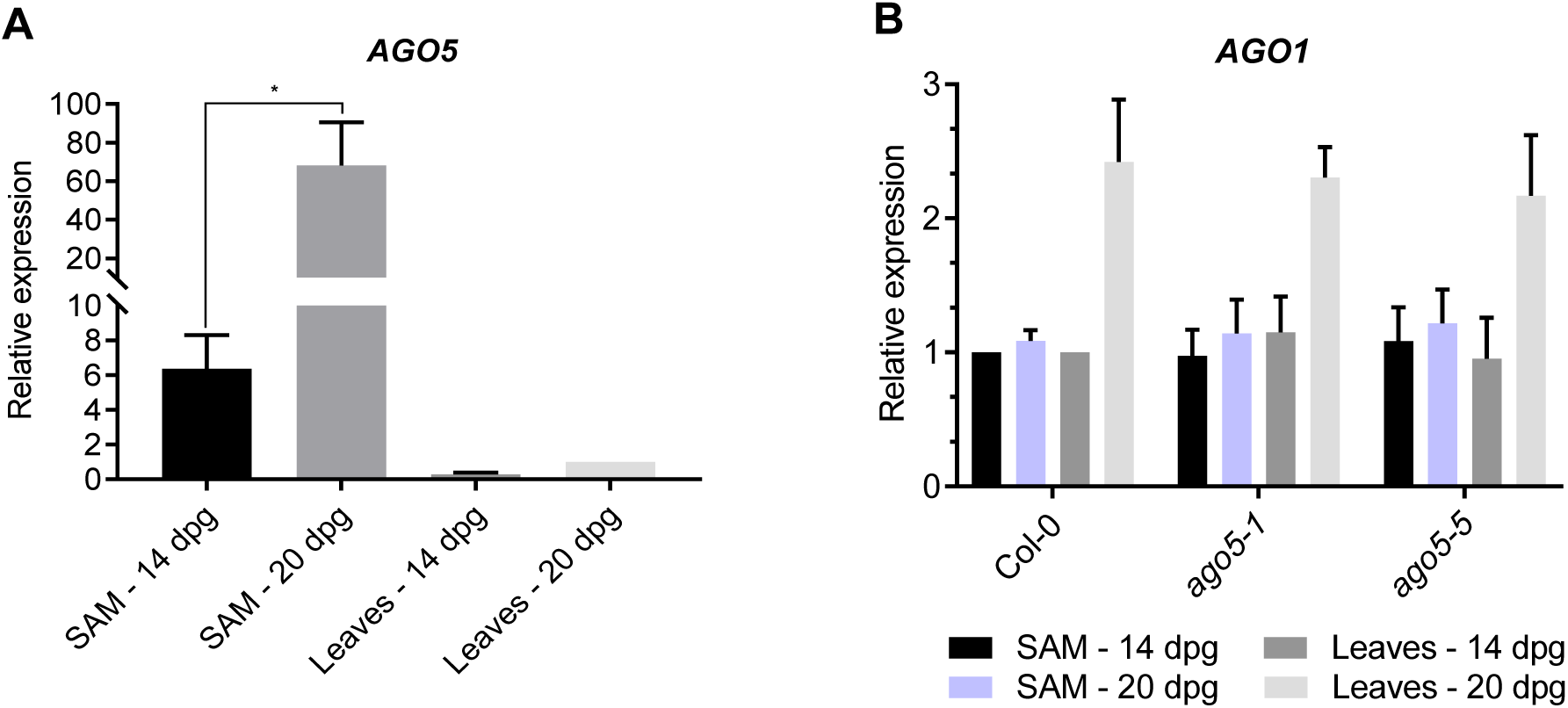
AGO5 is highly expressed in the shoot apical meristem. (**A**) RT-qPCR analysis of AGO5 gene expression in leaves and the SAM of Col-0. (**B**) RT-qPCR analysis of AGO1 gene expression in leaves and the SAM of Col-0 and ago5 mutants. Values are means ± SD of three biological replicates and are given as fold change compared to Col-0. Asterisks indicate significant differences from Col-0 in a Student’s t-test (*P < 0.05). Error bars indicate standard deviation.

AGO1 is an important player in developmental phase transition in plants and is presumed to have a similar miRNA binding capacity to AGO5, based on their miRNA sorting profiles (Mi et al., 2008). A potential crosstalk between the two AGO proteins is also supported by the expression profile of AGO1 and AGO5 during meristem development (Figure 2 – figure 1). Thus, we investigated *AGO1* expression in *ago5* mutants to confirm that AGO5 does not alter *AGO1* expression. Expression of *AGO1* was unaltered in both the SAM and the leaves of *ago5* mutants, as compared to wild-type plants (Figure 2B). Therefore, the early vegetative-to-reproductive phase transition of *ago5* mutants is unlikely to be caused by an altered expression of AGO1.

### Altered Expression of SQUAMOSA-PROMOTER BINDING LIKE Transcription Factors in *ago5* Mutants

SPL transcripts are tightly regulated by the miR156/157 aging pathway. Both SPLs and miR156 are expressed in the SAM and in leaves, where their abundances are negatively correlated (Schmid et al., 2003; Xu et al., 2016). Based on this information, we hypothesized that the role of AGO5 in SAM maturation and flowering is likely due to miR156-dependent targeting of SPL transcripts. To investigate this, we collected the SAMs of 20 dpg plants and quantified the relative abundance of six SPL gene transcripts by RT-qPCR in *ago5-1* and *ago5-5* mutants, as well as wild-type plants. Transcripts of SPL genes involved in meristem identity (SPL3, 4, 5), as well as shoot maturation and floral induction (SPL9, 11, 15) were all significantly more abundant in the SAMs of *ago5* mutants, as compared to Col-0 (Figure 3A, B). We also investigated the expression of the same genes at an earlier stage of development, when miR156 levels are higher. Significant differences in SPL expression levels were not apparent in *ago5* mutants relative to wild-type in the SAM of 14 dpg plants (Figure 3 – figure supplement 1). This suggests that the differences in SPL expression are not constitutive and that AGO5 is likely important for fine-tuning of developmentally regulated genes nearer to the reproductive stage.

**Figure 3.**
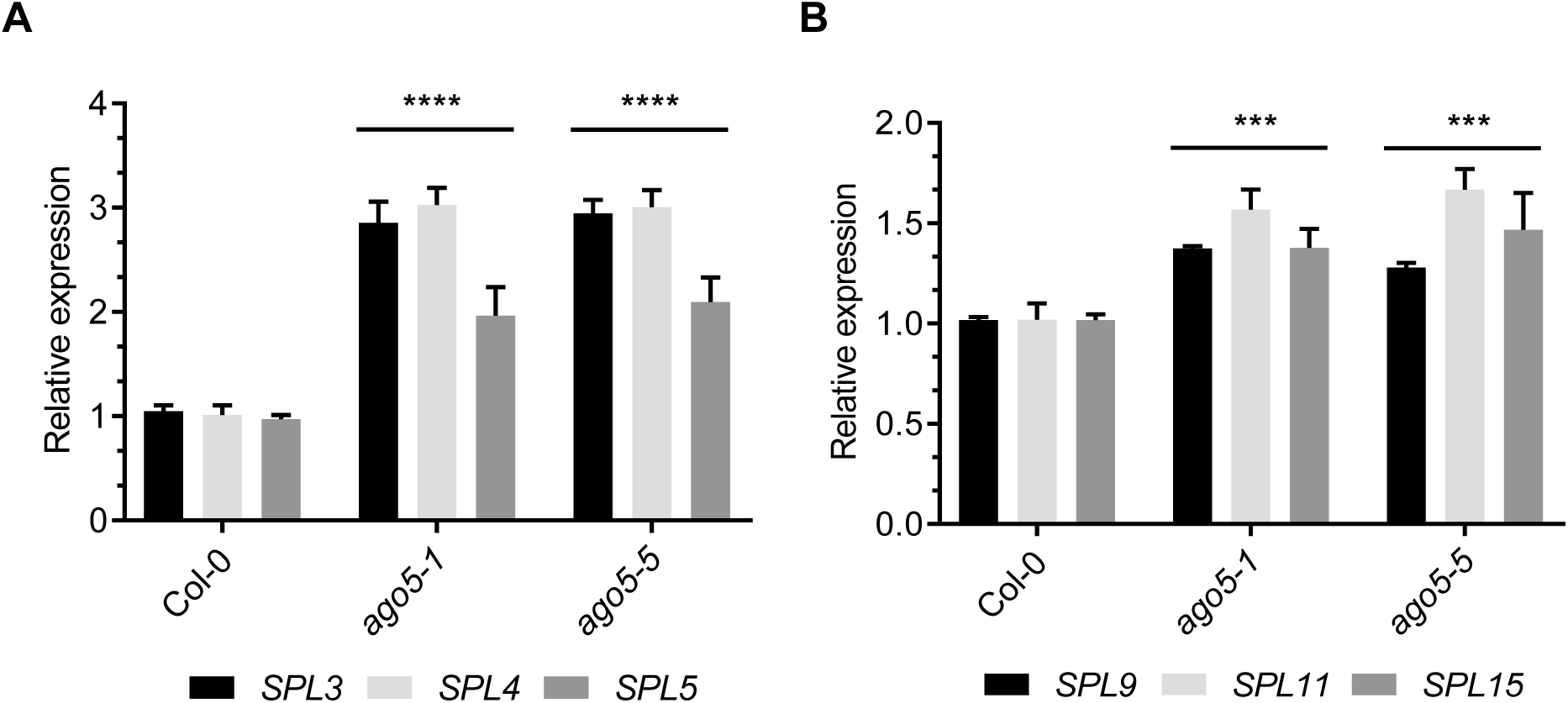
Altered expression of SPL transcription factors in *ago5* mutants. (**A**) RT-qPCR analysis of *SPL3/4/5* gene expression in the SAM of 20 days old Col-0, *ago5-1*, *ago5-5* plants. (**B**) RT-qPCR analysis of *SPL9/11/15* gene expression in the SAM of 20 days old Col-0, *ago5-1*, *ago5-5* plants. Values are means ± SD of three biological replicates and are given as fold change compared to Col-0. Asterisks indicate significant differences from Col-0 in a Student’s t-test (*P < 0.05, * * P < 0.01, * * * P < 0.001, * * * * P < 0.0001). Error bars indicate standard deviation.

Levels of pri-miR156 are reduced near the onset of reproductive phase change to allow the translation of developmentally regulated proteins (Xu et al. 2016). We thus explored whether pri-miR156 levels were different in 20 dpg *Arabidopsis* Col-0 and *ago5* mutants. Interestingly, we noticed that levels of pri-miR156 were approximately 35% lower in *ago5* mutants relatively to Col-0 (Figure 3 – figure supplement 2) suggesting that meristem identity shifts more rapidly from vegetative to reproductive phase in *ago5* mutants.

### ARGONAUTE5 Physically Interacts With miR156 *in planta*

In *Arabidopsis*, miRNAs with a 5’ terminal uridine, such as miR156, are preferentially loaded onto AGO1 (Mi et al. 2008). To investigate whether AGO5 and miR156 interact physically, we immunoprecipitated AGO5 from total extracts from one-week-old seedlings. Subsequent stem-loop amplification with miRNA156-specific primers indicated the presence of miR156 in immunoprecipitates from Col-0 and miR156 overexpressing plants but not the *ago5-5* mutant (Figure 4A). Surprisingly, miR156 levels in the input appeared to be lower in the *ago5-5* mutant, suggesting the miR156 is less stable in plants lacking AGO5.

**Figure 4.**
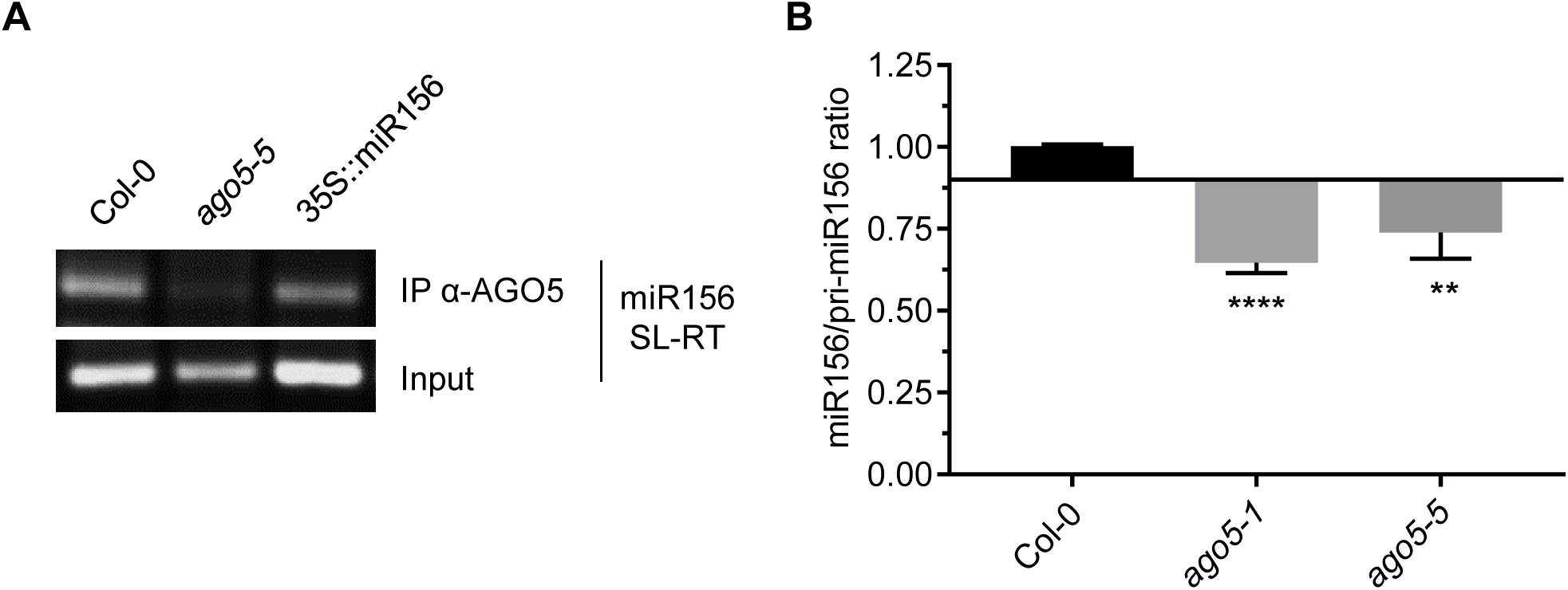
AGO5 binds and stabilizes miR156. (**A**) RNA extracted from AGO5 immunoprecipitates from one-week-old seedlings was subjected to stem-loop reverse transcription (SL-RT) using miR156-specific primers. (**B**) Ratios of miR156 to pri-miR156, based on Cq values obtained by RT-qPCR performed on RNA extracted from the SAMs of 20 day-old Col-0 and ago5 mutant plants, as indicated. Values are means ± SD of three biological replicates and are given as fold change compared to Col-0. Asterisks indicate significant differences from Col-0 in a Student’s t-test (*P < 0.05, * * P < 0.01, * * * P < 0.001, * * * * P < 0.0001). Error bars indicate standard deviation.

miRNAs are known to be more stable when they are bound by AGO proteins (Vaucheret et al. 2004). We thus quantified and compared the ratios of miR156 to pri-miR156 transcript levels in Col-0, *ago5-1* and *ago5-5* by qRT-PCR. In agreement with the above observation, we found a decrease in the ratio of miR156 to pri-miR156 in both *ago5* mutant lines compared to WT. These results suggest that miR156 is less stable in *ago5* compared to Col-0 (Figure 4B), suggesting that AGO5 may be the major binding partner for miR156 in the SAM.

### AGO5 is required for the Delay in Vegetative-to-Reproductive Phase Transition Induced by miR156 Overexpression

In wild-type *Arabidopsis* plants, constitutive overexpression of miR156 from the *CaMV 35S* promoter leads to a delay in juvenile-to-adult and vegetative-to-reproductive phase transition due to its negative regulation of SPL transcripts. To explore the role of AGO5 in the mir156-dependent regulation of SPL genes, we introduced a previously described 35S::miR156 transgene (Wu et al., 2009) into an *ago5-5* mutant background. These 35S::miR156/*ago5-5* plants displayed morphological characteristics similar to those of the 35S::miR156 alone, including altered leaf shape and stem length (Figure 5A). However, 35S::miR156*/ago5-5* plants bolted earlier than the 35S::miR156 under LD photoperiod, with flowering times not significantly different from Col-0 (Figure 5D). Thus, the effect of miR156 overexpression on bolting time is abrogated in the absence of AGO5. However, 35S::miR156*/ago5-5* plants did not bolt as early as *ago5-5* mutants, possibly because miR156 is able to load onto other AGO when artificially overexpressed.

**Figure 5.**
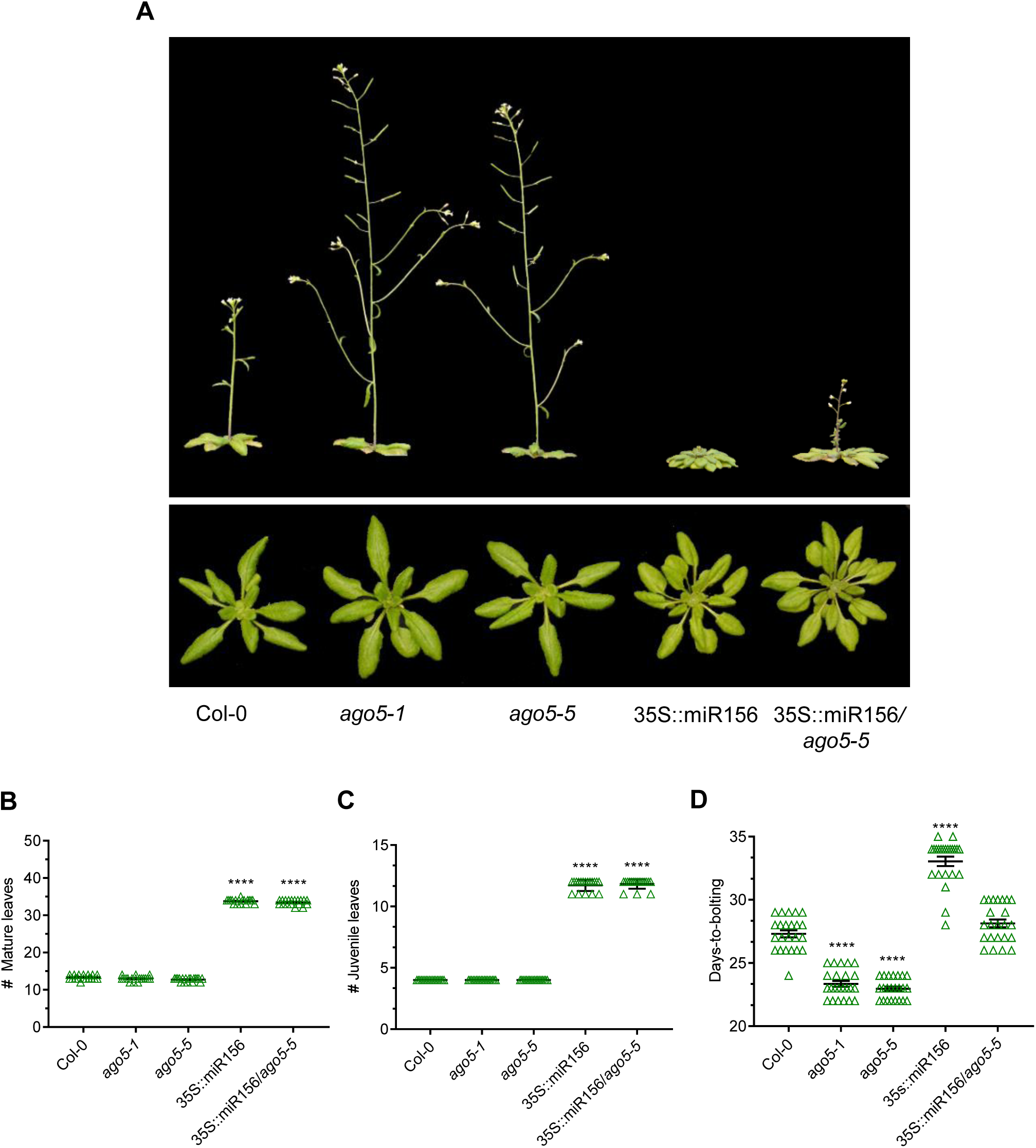
AGO5-miR156 interaction is crucial for a conventional regulation of vegetative-to-reproductive phase transition. Phenotypes of Col-0 and ago5 T-DNA insertion mutants (ago5-1 and ago5-5), 35S::miR156 and 35S::miR156/ago5-5. (**A**) Whole plant phenotypes. Photos were taken at 30 dpg. (**B**) Number of mature leaves (n = 23 plants). (**C**) Number of juvenile leaves (n = 23 plants). (**D**) Days-to-bolting (n = 23 plants). Experiments were repeated at least three times and similar results were obtained. Asterisks indicate significant differences from Col-0 in a Student’s t-test (*P < 0.05, * * P < 0.01, * * * P < 0.001, * * * * P < 0.0001). Error bars indicate standard deviation.

We characterized additional morphological aspects of the juvenile-to-adult phase transition in the 35S::miR156*/ago5-5* plants. Despite the marked difference in flowering times, no difference was observed between the 35S::miR156 and the 35S::miR156/*ago5-5* lines with respect to either aerial architecture (Figure 5A) or mature and juvenile leaf counts (Figure 5B, C). These observations are consistent with the fact that *AGO5* expression in the leaves is virtually undetectable (Figure 5 – figure supplement 1). Thus, miR156 may act through AGO5 in meristems, but in leaves its effects may be mediated through other AGO proteins, such as AGO1, which has previously been shown to modulate juvenile-to-adult phase transition in leaves.

### AGO5 Is Essential for a Conventional Regulation of the miR156 Aging Pathway in the SAM

To support a functional relationship at the molecular level between AGO5 and mir156 in regulating vegetative-to-reproductive transition, we compared expression levels of SPL genes and developmentally regulated genes (*AP1*, *SOC1*, *FUL and* LFY) at 20 dpg in Col-0, *ago5-1*, *ago5-5*, 35S::miR156 and 35S::miR156*/ago5-5*. As expected, *ago5* mutants showed higher levels of expression of all SPL genes (Figure 6A), as well as SPL-regulated genes (Figure 7A), compared to all other genotypes. In contrast, both 35S::miR156*/ago5-5* and 35S::miR156 plants, which have not bolted at this timepoint, showed dramatically reduced expression levels of the same genes compared to Col-0 at 20 dpg (Figure 6A, 7A).

**Figure 6.**
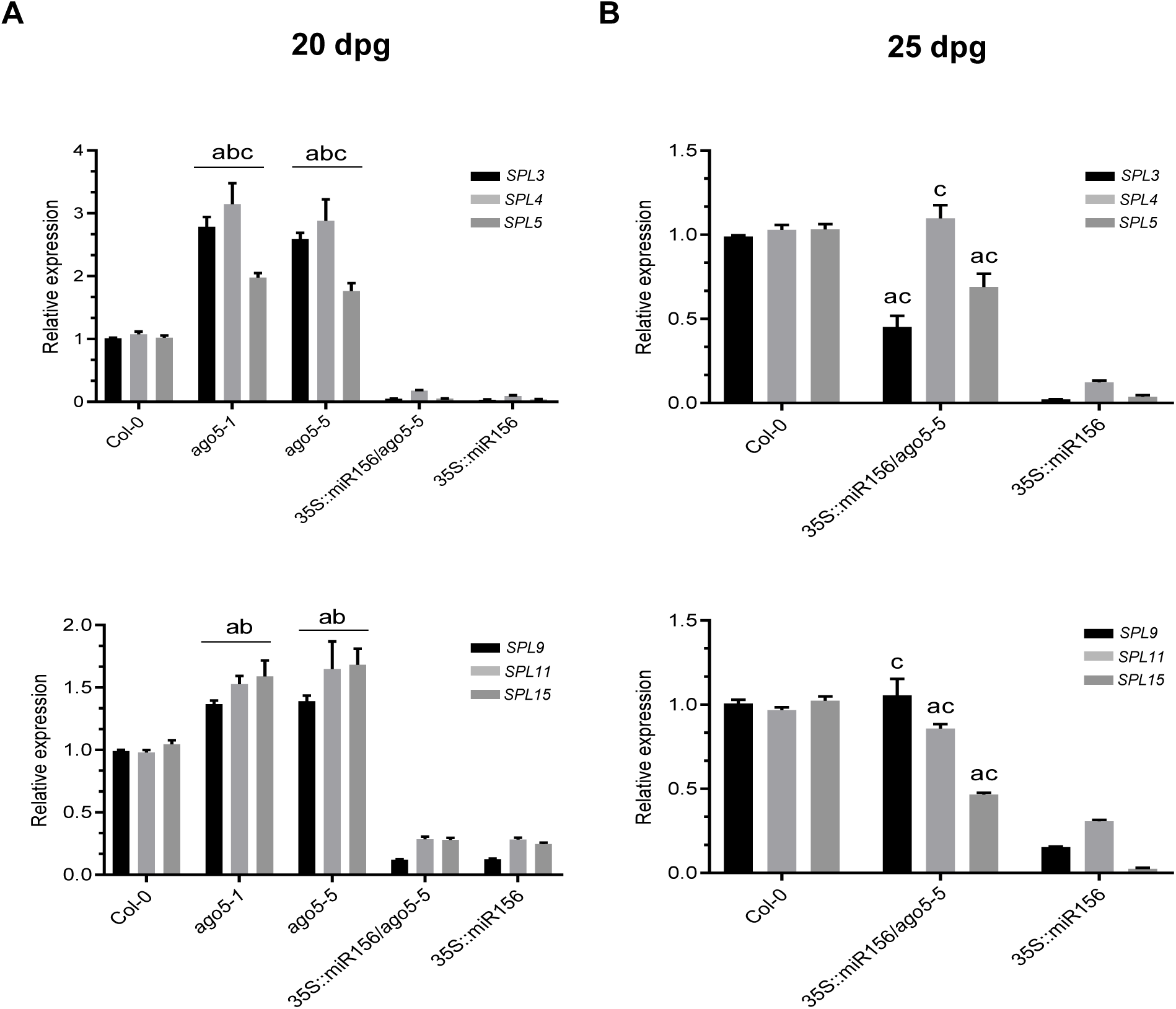
*AGO5* regulates fine-tuning of SPL transcription factor expression. (**A**) RT-qPCR analysis of SPL gene expression in the SAM of 20 dpg Col-0, *ago5-1*, *ago5-5*, 35S::miR156 and 35S::miR156/*ago5-5* plants. (**B**) RT-qPCR analysis of SPL gene expression in the SAM of Col-0, 35S::miR156 and 35S::miR156/*ago5-5* of 25 dpg plants. Values are means ± SD of three biological replicates and are given as fold change compared to Col-0. a = significantly different from Col-0; b = significantly different from 35S::miR156/*ago5-5*; c = significantly different from 35S::miR156 (Student’s t-test; P < 0.01). Error bars indicate standard deviation.

**Figure 7.**
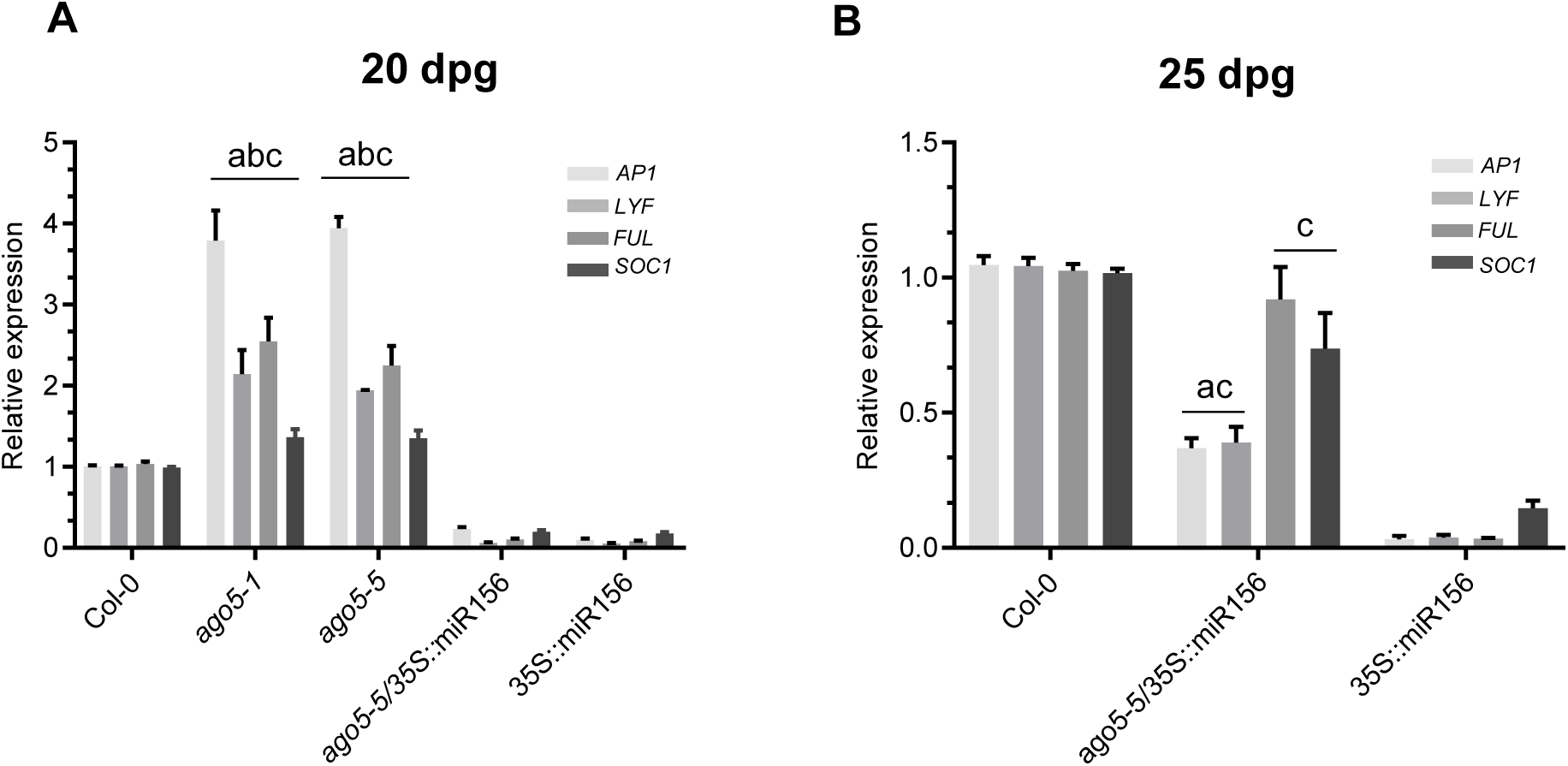
Developmentally regulated gene expression in the 35S::miR156/ago5-5 hybrid. (**A**) RT-qPCR analysis at 20 dpg of *AP1*, *LYF*, *FUL* and *SOC1* gene expression in the SAMs Col-0, *ago5-1*, *ago5-5*, 35S::miR156 and 35S::miR156/*ago5-5* plants. (**B**) RT-qPCR analysis at 25 dpg of *AP1*, *LYF*, *FUL* and *SOC1* gene expression in the SAMs of Col-0, 35S::miR156 and 35S::miR156/*ago5-5* plants. Values are means ± SD of three biological replicates and are given as fold change compared to Col-0. a = significantly different from Col-0; b = significantly different from 35S::miR156/*ago5-5*; c = significantly different from 35S::miR156 (Student’s t-test; P < 0.01). Error bars indicate standard deviation.

Sinc*e* 35S::miR156*/ago5-5* plants bolt at the same time as Col-0 plants, we evaluated the expression of SPL genes near the Col-0 bolting time (*ago5* mutants were not included as these plants have already bolted at this time point). As expected, we found that expression levels of all of SPL genes analyzed in 35S::miR156*/ago5-5*, but not 35S::miR156 plants, were close, or equal to wild type at 25 dpg (Figure 6B). Expression levels of developmentally regulated gene was also re-established in *35S::miR156/ago5-5* compared to 35S::miR156 plants (Figure 7B). As such, these results further indicate a role for AGO5 in the fine-tuning of gene regulation during the vegetative-to-reproductive phase transition.

## DISCUSSION

Aging pathway studies have shown important roles for microRNAs in floral induction, in particular miR156. The latter is known to regulate this pathway by targeting SPL transcription factors, which are in turn responsible for the transcription of genes involved in the appearance of adult and reproductive traits (J.-W. Wang, 2014). To affect gene expression, miRNAs must act through AGO proteins and our data suggest that AGO5 plays an important role in the miR156 aging pathway in meristematic tissues.

We observed an early developmental phase transition phenotype in *ago5* mutant plants, supported by precocious SPL expression. Furthermore, we found that AGO5 was able to bind to miR156 in seedlings and this miRNA was less stable in the SAM of *ago5* mutants compared to wild-type plants. In agreement with this, interactions between AGO5 and miR156 have also reported in inflorescent tissues of *Arabidopsis* in large-scale AGO-miRNA interaction studies (Mi et al., 2008; Takeda, Iwasaki, Watanabe, Utsumi, & Watanabe, 2008; Shao et al., 2014). In support of a major role of AGO5 in the vegetative-to-reproductive phase transition, we found that the delay in bolting time was abolished in miR156 overexpressing plants lacking *AGO5*. Thus, this suggests that the late flowering phenotype of miR156-overexpressing plants is due mainly to its interaction with AGO5. However, we do not rule out possible compensation by another AGO protein in the presence of artificial overexpression of miR156 in these lines.

AGO5 appears to be involved in several plant development processes including seed pigmentation in *Glycine max* (Cho, Jones, & Vodkin, 2017), sporogenesis in *Oryza sativa* (Nonomura et al., 2007), establishment of nodules in legume-rhizobia interactions (Reyero-Saavedra et al., 2017), as well as megagametogenesis (Tucker et al., 2012) and pollen development in *Arabidopsis* (Borges, Pereira, Slotkin, Martienssen, & Becker, 2011). In *Arabidopsis*, *AGO5* is highly and almost exclusively expressed during all stages of flowers and seed formation (Kapoor et al., 2008; Schmid et al., 2005; Waese et al., 2017), including inflorescence meristems (Mantegazza et al., 2014), suggesting a primordial role for this protein in the reproductive phase of development. We find that *AGO5* is also highly expressed in the SAM of adult plants where it appears to target SPL transcripts in a miR156-dependent manner. At the same time, AGO1 has previously identified as an important player in plant development (Baumberger & Baulcombe, 2005; Vaucheret et al., 2004). Supporting this, AGO1 binds miR156 and miR157 in *Arabidopsis* seedlings, suggesting a role for AGO1 in the vegetative phase change (He et al., 2018; Azevedo et al. 2010). However, a role for AGO1 in miR156-dependant regulation of flowering time has not been demonstrated, possibly in part because *Arabidopsis ago1* mutant phenotypes can be difficult to interpret due to dramatic pleiotropic developmental defects (Vaucheret et al. 2004). In our analysis, *AGO1* shows a ubiquitous expression, as opposed to the tissue-specific expression of *AGO5*, which is restricted mainly to the SAM and reproductive tissues (Figure 2 – figure supplement 1). The phenotypic aspect of 35S::miR156/*ago5-5* plants suggest that AGO1 and AGO5 might both act in the miR156 aging pathway but that they may have distinct functions in different tissues. While these plants displayed a flowering time comparable to wild-type plants, the leaf architecture resembled that of the miR156-overexpressing plants. Since miR156 is also known to be important for leaf development and morphology (Feng et al., 2016; Xie et al., 2012), we suggest that the miR156-dependant regulation of leaf development is likely due to its interaction with another AGO protein, most probably AGO1. This would be consistent with the near absence of *AGO5* expression in leaves and the absence of a leaf morphology phenotype in both *ago5* mutants.

In later stages of plant development, other factors promote the expression of SPL genes above the threshold set by miR156, enabling phase transitions to occur. Expression levels of *SPL3/4/5* are influenced by miR156-independent floral inductive signals such as CONSTANS (CO) and FLOWERING LOCUS T (FT) (Jung, Ju, Seo, Lee, & Park, 2012). A crosstalk between Gibberellic acid (GA) and miR156-regulated pathways has also been reported. In both long and short day photoperiods, GA has been shown to play an important role in the expression of SPLs and in their targets, such as SOC1 and FUL (Galvao, Horrer, Kuttner, & Schmid, 2012; Moon et al., 2003; Porri, Torti, Romera-Branchat, & Coupland, 2012). Interestingly, miR156 levels are largely unaffected by the activity of the aforementioned pathways. Despite low levels of miR156 in late developmental stages, He et al. (2018) suggested that this miRNA may act in fine-tuning SPL expression in a threshold-dependent manner. The expression of *AGO5* specifically in the SAM and the higher abundance of SPL transcripts in *ago5* mutants near bolting time suggest that AGO5 plays a role in miR156-mediated vegetative-to-reproductive phase transition by acting on meristem identity. In agreement with this, when the expression of SPL and SPL-regulated genes was assessed in Col-0, 35S::miR156 and the 35S::miR156*/ago5-5* at a later stage of the vegetative-to-reproductive phase change, we observed similar expression patterns between Col-0 and the 35S::miR156*/ago5-5*. In contrast, in 20 day-old *35S::miR156/ago5-5* plants, no significant difference was observed with respect to SPL and SPL-regulated gene expression as compared to the *35S::miR156* line alone. This observation reinforces the hypothesis that AGO5 mediates fine-tuning of SPLs through its interaction with the miR156 in the SAM.

To conclude, we propose a model whereby a novel AGO-miRNA interaction mediates fine-tuning of vegetative-to-reproductive phase transition in *Arabidopsis*. Through genetic and biochemical approaches, we confirm that a physical and functional interaction between AGO5 and miR156 is necessary for regulating the miR156-aging pathway in the SAM but not in leaves. As such, this study highlights the complexity by which the RNA silencing machinery modulates different aspects of plant development by allowing different AGO proteins to act in tissue-specific manners.

## MATERIALS AND METHODS

### Plant Material and Growth Conditions

All *Arabidopsis thaliana* plants were grown in Agromix (Fafard) in growth chambers with 16h-light and 8h dark photoperiod at 22°C. Except for the 35S::miR156/*ago5-5* hybrid, all *Arabidopsis* mutant lines have been previously described including *ago5-1* (Katiyar-Agarwal, Gao, Vivian-Smith, & Jin, 2007), *ago5-5* (Brosseau and Moffett., 2015) and 35S::miR156 (Wu et al., 2009). The 35S::miR156/*ago5-5* line was generated by crossing the *35S:miR156* transgenic line with *ago5-5* mutant. Newly generated lines were genotyped using the primers listed in Supplementary Table 1.

### RNA Extraction and Treatment

Total RNA was extracted using Trizol (Invitrogen) and then treated with RNase-free DNase (Ambion) following the manufacturer’s instructions. Meristem tissue was harvested as described (Xu et al. 2016). The quantity and purity of total RNA samples were determined from 280/260 and 260/230 nm absorbance ratios measured using a Nanophotometer MBI Lab Equipment. Ratios of 1.8 and 2.3, respectively, were accepted as pure RNA and electrophoresis with agarose gel 2% was carried out to observe RNA integrity. Following RNA extraction, 2 μg of RNA was reverse transcribed with SuperScript II (Invitrogen) according to the manufacturer’s instructions.

### RT-qPCR and Gene Expression Analysis

RT-qPCR analysis was performed using a Biorad CFX96 real-time PCR system (Bio-Rad) operated by CFX Manager software (version 3.0). In all assays, the samples analyzed included technical and biological triplicates. Each biological replicate included meristems from eight different plants. The specificity of each primer pair was verified from the dissociation (melt) curves acquired after 40 cycles using CFX Manager software (version 3.0). PCR amplification efficiency (E) was determined for the validation of primers according to the standard curve method after using a set of all cDNA samples with 5-fold serial dilution. The RT-qPCR was programmed with an initial step of 98°C for 2 min and 40 cycles of 98°C for 2 s and annealing/extension at 60°C for 5 s, with melt curve analyses from 65 to 95°C in 0.5°C increments. *ACTIN2* was used as a reference gene for the normalization of the relative expression data as described previously (Xu et al., 2016). Average Cq values were then normalized by ΔΔCT formula against the indicated reference gene. The specific primer efficiency was considered in the relative gene expression analysis as suggested previously (Pfaffl, 2001). Primer sequences used in this study and their efficiency are listed in Supplementary Table 1.

### miRNA Immunoprecipitation

AGO-RNA complexes were obtained by grinding one-week-old plants with a mortar in cold extraction buffer and immunoprecipitated (IP) as described previously (Brosseau & Moffett, 2015). For input, small RNAs were obtained from 100 μl of AGO-RNA grounded samples and extracted as previously described (Varkonyi-Gasic et al.). RNA was extracted from AGO IP complexes using Trizol (Invitrogen). Thereafter, miRNA reverse transcription and amplification was performed as described, with modifications (Varkonyi-Gasic et al.). PCR was performed using Taq polymerase and cycles were as followed (94°C for 2 minutes; 94°C for 15 seconds; 53°C for 25 seconds; 72°C for 20 seconds; 50 cycles).

### Microscopy Assay

Floral architecture was verified under a Leica M165FC stereomicroscope with a LeicaM170HD camera. All photos were taken with a 0.63X objective.

## ACKNOWLEDGMENTS

This study was supported by a Discovery grant from the Natural Sciences and Engineering Research Council (Canada) to P.M. and by fellowships to C.R-L. and G. S-M from the Agrophytosciences CREATE program. We are thankful to Daniel Garneau for help with stereomicroscopy and Marie Bernadette Dib Diam for providing the 5.8S rRNA primer.

---

**Figure 1 - figure supplement 1.**
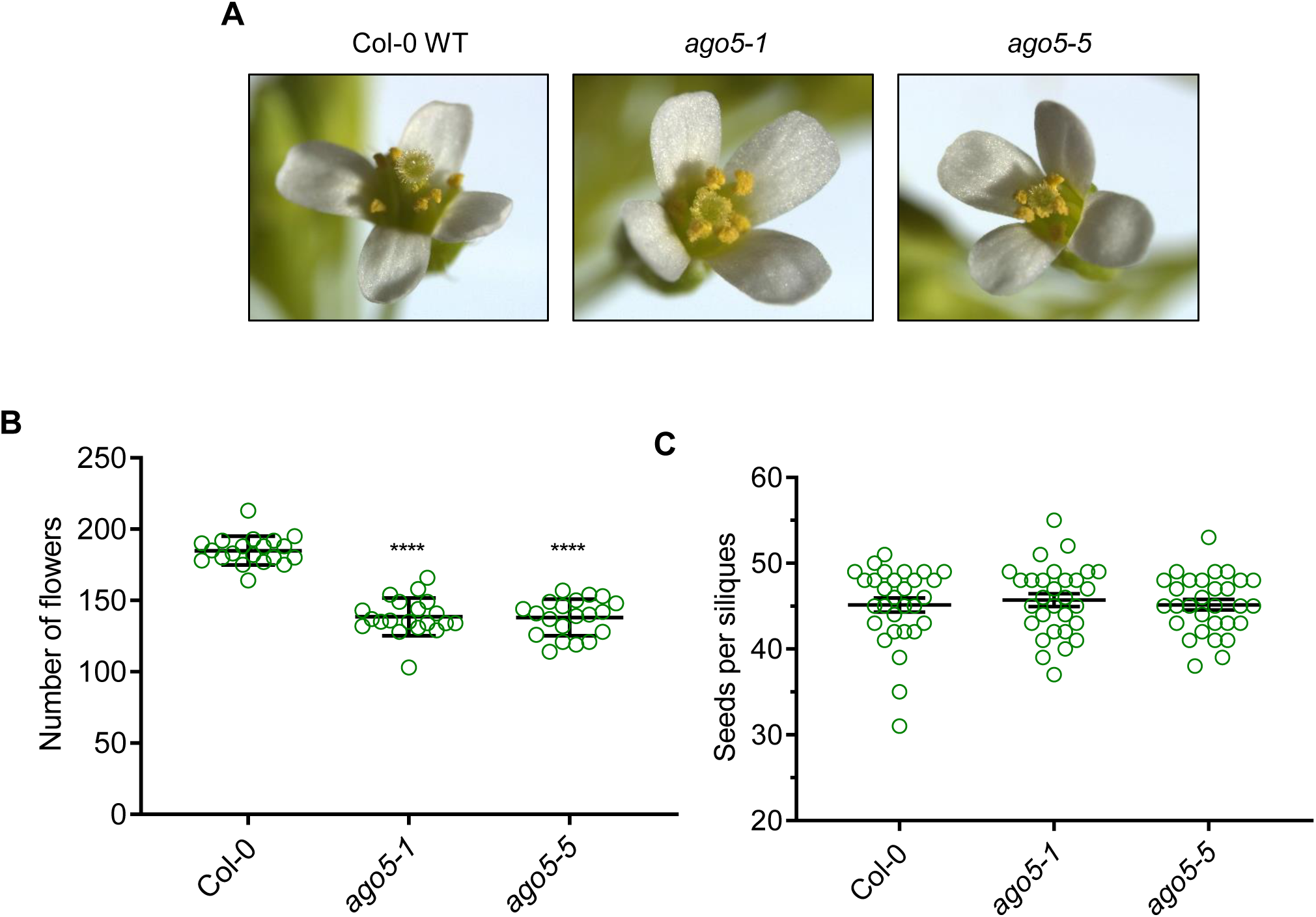
*AGO5* is important for floral output, but not architecture. (**A**). Phenotypes of Col-0 and *ago5* T-DNA insertion mutant (*ago5-1* and *ago5-5*) flowers. (**B**) Number of flowers present on individual plants, per genotype (n = 26). (**C**) Number of seeds per silique (n = 98). Values are means ± SEM of three biological replicates and are given as fold change compared to Col-0. Asterisks indicate significant differences from Col-0 in a Student’s t-test (*P < 0.05, * * P < 0.01, * * * P < 0.001, * * * * P < 0.0001). Error bars indicate standard deviation.

**Figure 2 - figure supplement 1.**
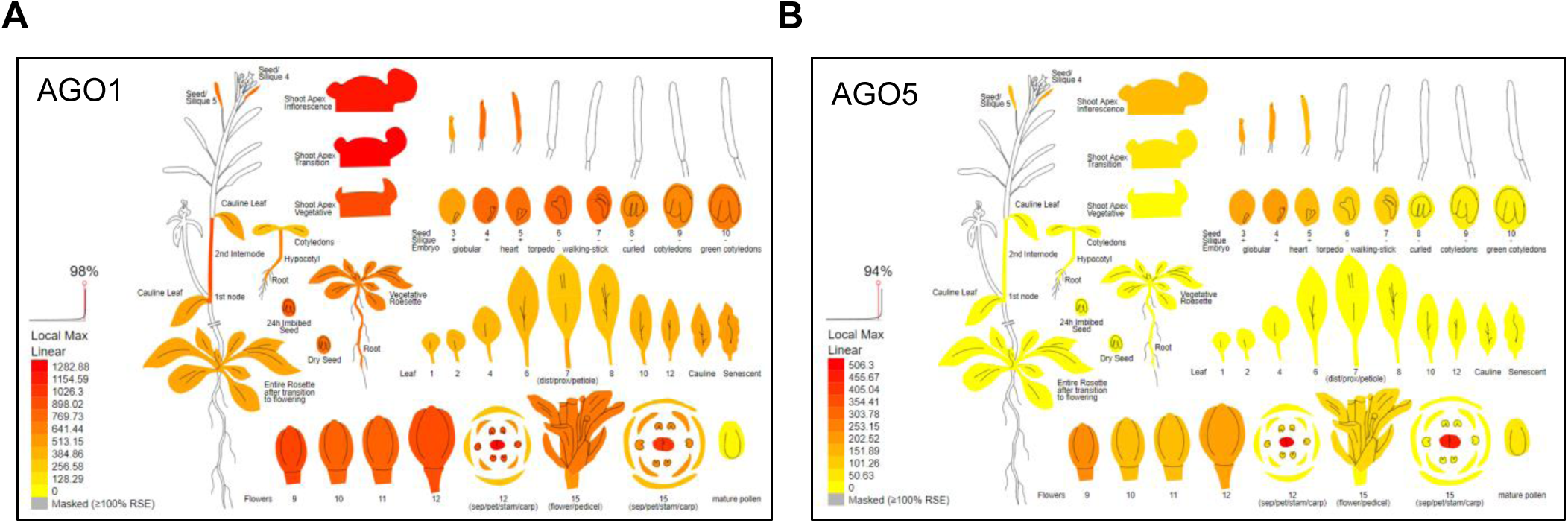
Expression profiles of *AGO1* and *AGO5* in *Arabidopsis thaliana*. **A.** Heatmap of *AGO1* expression levels in diverse life stages and tissues of *Arabidopsis thaliana*. **B.** Heatmap of *AGO5* expression levels in diverse life stages and tissues of *Arabidopsis thaliana*. Data obtained from http://bar.utoronto.ca/eplant/ (Waese et *al*. 2017).

**Figure 3 - figure supplement 1.**
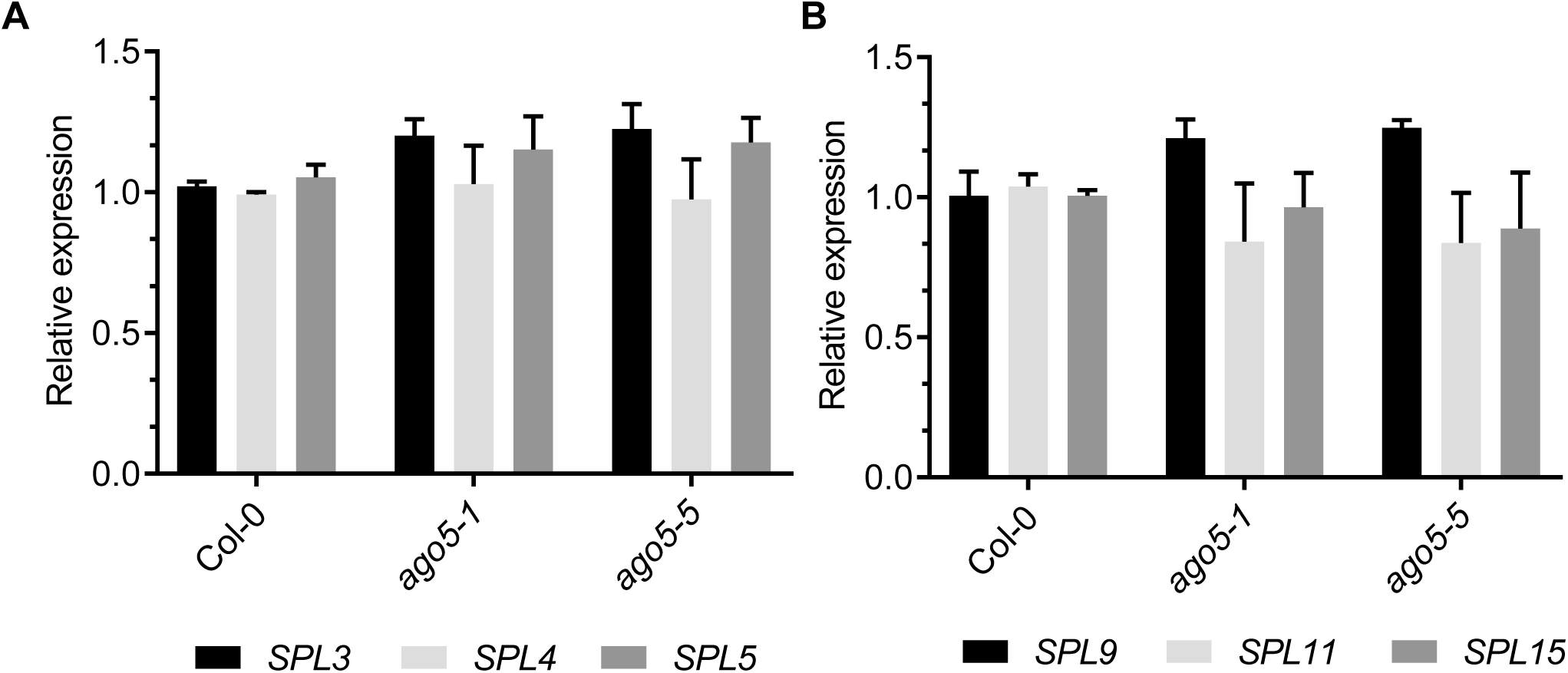
SPL expression levels in the SAM of two-week old *ago5* mutants. (**A**) RT-qPCR analysis of *SPL3/4/5* gene expression in the SAM of 14-day-old Col-0, *ago5-1*, *ago5-5* plants. (**B**) RT-qPCR analysis of *SPL9/11/15* gene expression in the SAM of 14-day-old Col-0, *ago5-1*, *ago5-5* plants. Values are means ± SD of three biological replicates and are given as fold change compared to Col-0. Asterisks indicate significant differences from Col-0 in a Student’s t-test (*P < 0.05, * * P < 0.01, * * * P < 0.001, * * * * P < 0.0001). Error bars indicate standard deviation.

**Figure 3 - figure supplement 2.**
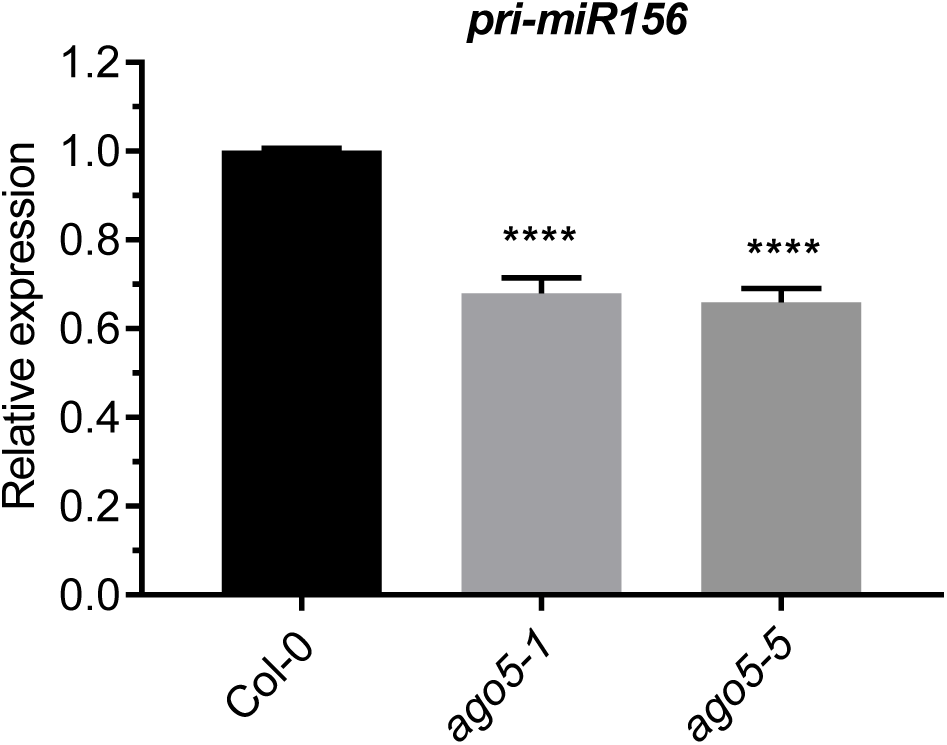
Lower levels of *pri-miR156* in the SAM of *ago5* mutants near bolting time. RT-qPCR analysis of pri-miR156 gene expression in the SAM of 20-day-old Col-0, *ago5-1* and *ago5-5* plants. Values are means ± SD of three biological replicates and are given as fold change compared to Col-0. Asterisks indicate significant differences from Col-0 in a Student’s t-test (*P < 0.05, * * P < 0.01, * * P < 0.001, * * * * P < 0.0001). Error bars indicate standard deviation.

**Figure 5 - figure supplement 1.**
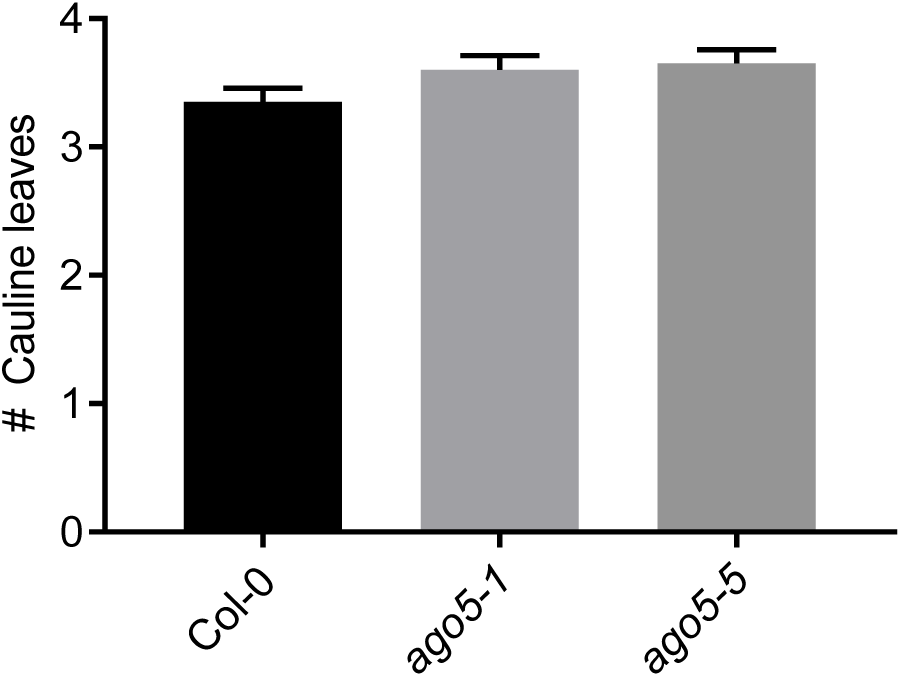
Cauline leaf numbers are unaltered in *ago5* mutants. Number of cauline leaves in Col-0, *ago5-1* and *ago5-5* plants. Values are means ± SD of three biological replicates and are given as fold change compared to Col-0. Error bars indicate standard deviation.

**Figure 5 - figure supplement 2.**
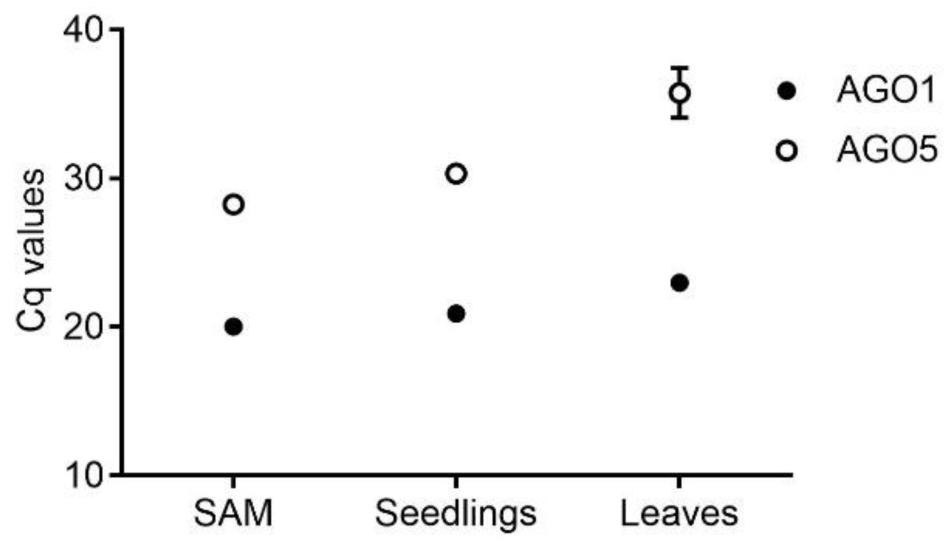
Cycle of quantification (Cq) values of *AGO1* and *AGO5* in *Arabidopsis thaliana* in one-week-old seedlings and 20-day-old SAM and leaf tissue. Values are means ± SD of three biological replicates.

**Supplementary Table 1.**
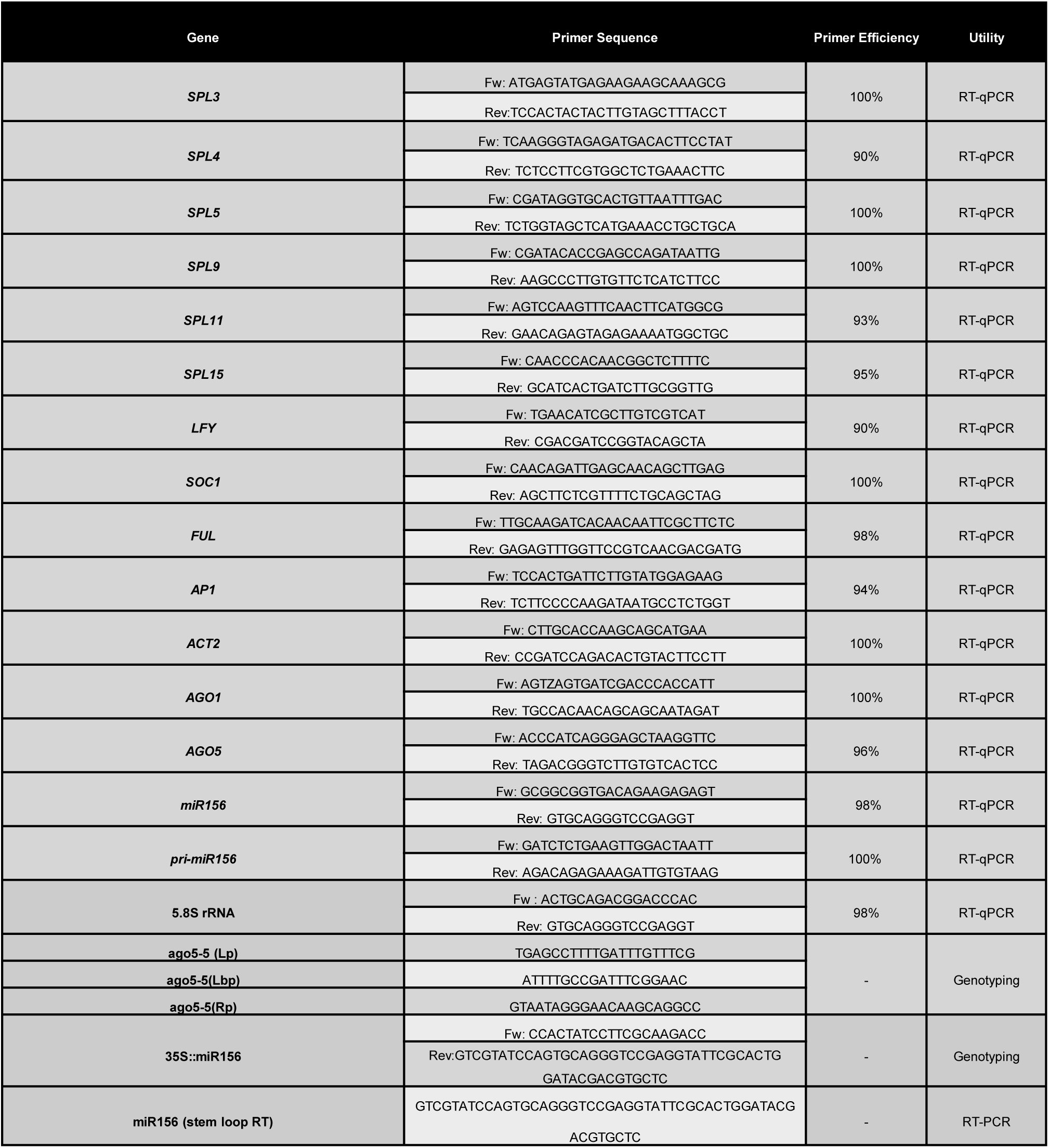
Oligos used in this study.

